# Isolation of high yield and quality RNA from human precision-cut lung slices for RNA-sequencing and computational integration with larger patient cohorts

**DOI:** 10.1101/2020.08.21.254516

**Authors:** John Stegmayr, Hani N. Alsafadi, Wojciech Langwiński, Anna Niroomand, Sandra Lindstedt, Nicholas D. Leigh, Darcy E. Wagner

## Abstract

Precision-cut lung slices (PCLS) have gained increasing interest as a model to study lung biology and disease, as well as for screening novel therapeutics. In particular, PCLS derived from human tissue can better recapitulate some aspects of lung biology and disease as compared to PCLS derived from animals (*e.g.* clinical heterogeneity), but access to human tissue is limited. A number of different experimental readouts have been established for use with PCLS, but obtaining high yield and quality RNA for downstream gene expression analysis has remained challenging. This is particularly problematic for utilizing the power of next-generation sequencing techniques, such as RNA-sequencing (RNA-seq), for non-biased and high through-put analysis of PCLS human cohorts. In the current study, we present a novel approach for isolating high quality RNA from a small amount of tissue, including diseased human tissue, such as idiopathic pulmonary fibrosis (IPF). We show that the RNA isolated using this method is of sufficient quality for both RT-qPCR and RNA-seq analysis. Furthermore, the RNA-seq data from human PCLS was comparable to data generated from native tissue and could be used in several established computational pipelines, including deconvolution of bulk RNA-seq data using publicly available single-cell RNA-seq data sets. Deconvolution using Bisque revealed a diversity of cell populations in human PCLS derived from distal lung tissue, including several immune cell populations, which correlated with cell populations known to be present and aberrant in human disease, such as IPF.

## Introduction

Precision-cut lung slices (PCLS) have received increased attention as a relevant model for studying and modeling lung diseases (1). Most lung diseases involve several cell types and derangements in their surrounding environment, such as the extracellular matrix. They thus require complex disease models to explore disease pathogenesis and novel therapeutics. PCLS excel at retaining the native architecture of the lung and keeping the spatial composition of cells and their surrounding environment intact, making it possible to study cell-cell and cell-matrix interactions in health or disease (1).

While cell-based screening assays have been invaluable at measuring drug cytotoxicity and assessing potential therapeutic effects, they remain limited in predicting clinical efficacy due to a lack of the complex cellular and environmental contexts in which a disease develops (2). The high failure rate of pre-clinically selected drugs, when tested at later clinical stages, has been attributed to the inherent lack of complexity in conventional drug screening and validation. Subsequently, use of more advanced models for drug screening, such as 3D engineered hydrogels and lung-on-a-chip, are emerging as screening tools for drug candidates (3, 4). Yet these models are limited by the source of materials used to generate the disease model such as the cell type or extracellular matrix component used. The complexity of PCLS and their ability to recapitulate the native environment of the lung makes it well-suited for advanced drug screening (5). Additionally, PCLS can provide readouts from multiple levels, such as gene/protein expression and imaging, which allows for both phenotypic and genotypic profiling after exposure to a specific drug, including small molecules and biologics (1).

Gene expression analysis from PCLS has been previously performed by us and others, but has so far mainly been limited to techniques such as RT-qPCR. Next-generation sequencing, such as RNA-sequencing (RNA-seq), could be used for non-biased and high through-put analysis of transcriptional changes in PCLS. Furthermore, RNA-seq of PCLS would allow for integration of data with larger cohorts or types of genome-wide data (*e.g.* deconvolution using single-cell RNA-seq (scRNA-seq) data), functional profiling (*e.g.* pathway analysis), and much more (6, 7). However, current nucleic acid isolation techniques used for PCLS yield low and unreliable amounts at low quality. Additionally, the low yield of nucleic acids confers an additional limitation on the number of experimental conditions which can be examined in a single biological individual, as more PCLS tissue is needed for isolation. The high ratio of agarose to tissue in PCLS is thought to be the major factor for this limitation (8, 9), but thus far, techniques which can selectively eliminate agarose from PCLS without degrading nucleic acids have not been described.

In this study, we established a new RNA isolation protocol generating high yields of nucleic acids with sufficient quality for RNA-seq analysis from a small amount of PCLS, including normal and diseased human as well as animal lung tissue. Furthermore, we show that the generated RNA-seq data from human PCLS is of sufficient quality for downstream deconvolution analysis and integration with larger patient cohorts.

## Materials and methods

### Animal lung tissue

The use of mouse and pig lung tissue was approved by the Local Ethics Committee for Animal Research in Lund, Sweden (Dnr 6436/2017 and 8401/2017). All animal experiments were conducted in accordance with legislation by the European Union (2010/63/EU) and local Swedish law. All animals received care according to the Guide for the Care and Use of Laboratory Animals: Eighth Edition, National Academies Press (2011).

### Human lung tissue

The use of human lung tissue was approved by the Local Ethics Committee in Lund (Dnr 2017/396 and 2018/386). All experiments were conducted in accordance with the Declaration of Helsinki with written informed consent from all patients.

### Generation of mouse, pig, and human PCLS

For the generation of mouse PCLS, a C57BL/6NRj female mouse (~ 12 weeks old) (Janvier Labs) was sacrificed by an intraperitoneal injected overdose of ketamine and xylazine and the lungs were filled with agarose *in situ* and dissected as previously described (10). For the generation of pig and human PCLS, the lungs were filled with agarose *ex situ* as previously described for human lung tissue (10, 11). Porcine lung tissue was obtained from a healthy 4-month old pig (~ 60 kg) following 3 hours of mechanical ventilation at tidal volumes of 6 ml/kg with 5 cm H_2_O positive end expiratory pressure (PEEP) and FiO_2_ of 50 %. The pig was heparinized and euthanized by induction of ventricular fibrillation. Human PCLS were generated from lung explants from two patients: a 60-year-old female (non-smoker) undergoing size-mismatch surgery about 2 weeks after a lung transplantation (Patient 1) and a 54-year-old male (smoker, 20 pack/year) diagnosed with idiopathic pulmonary fibrosis (IPF) (Patient 2). For all species, 3% (weight/volume) low-gelling agarose (A9414, Sigma-Aldrich) in Dulbecco’s Modified Eagle Medium/Nutrient Mixture F-12 (DMEM/F-12) kept at 42 °C was used for lung filling. Before lung tissue slicing, the individual lobes of the mouse lung were separated and for the pig and human lungs, distal tissue was dissected into cubes (1.5-2.5 cm in all dimensions) and stored in DMEM/F-12 at 4 °C until slicing the same day. A 7000SMZ-2 Vibrotome (Campden Instruments Ltd.) equipped with a temperature controlled tissue bath (Campden Instruments Ltd.) containing DMEM/F-12 supplemented with 10 mM HEPES at 4-7 °C was used for tissue slicing. Mouse lung tissue was sliced at a thickness of 500 µm with stainless steel microtome blades at a frequency of 60 Hz and an amplitude of 1 mm, whereas pig and human tissue were sliced at a frequency of 90 Hz and an amplitude of 1.5 mm at a thickness of 500 µm. To normalize the size of the slices, a 4-mm diameter biopsy puncher was used. After slicing, the PCLS were transferred to DMEM/F-12 supplemented with 0.1 % fetal bovine serum, 100 U/ml penicillin, 100 µg/ml streptomycin, and 2.5 µg/ml amphotericin B and kept in a humidified incubator at 37 °C supplemented with 5 % CO_2_ in air until sample collection (2 hours for Patient 1 and 48 hours for Patient 2). Samples for RNA isolation were collected by flash freezing in liquid nitrogen and stored at −80 °C.

### RNA isolation

Three protocols were used; a standard protocol and two with further modifications. For all protocols, four 4-mm PCLS punches were used per isolation. The PCLS were homogenized using a Qiagen TissueLyser II with 5-mm stainless steel beads in the RNeasy Micro Kit provided lysis buffer (*i.e.* Buffer RLT supplemented with 143 mM 2-mercaptoethanol) (for standard protocol) or in TRIzol (for modified protocols 1 and 2) at 27 Hz for 3 × 1 minute. For the standard protocol, the lysis buffer homogenates were centrifuged (16,000 × g) for 3 minutes, whereupon the supernatants were mixed with an equal volume of 70 % ethanol and loaded onto a RNeasy MinElute Spin Column. For the second protocol (modified protocol 1), the TRIzol homogenates were mixed with chloroform 5:1 (volume/volume), incubated for 15 minutes, and were then centrifuged (12,000 × g) for 15 minutes at 4 °C. Following the centrifugation, the aqueous phase was mixed with an equal part 70 % ethanol and loaded onto a RNeasy MinElute Spin Column. For the third protocol (modified protocol 2), the TRIzol homogenates were mixed with 100 % ethanol at 2:1 (volume/volume) and loaded onto a RNeasy MinElute Spin Column. The following steps for the RNA enrichment were performed according to the instructions provided for the RNeasy Micro Kit, for all three protocols, with the inclusion of the optional DNase I (Qiagen) step for elimination of genomic DNA. The RNA was eluted with 14 or 20 µl water and stored at −80 °C until further analysis. RNA from native pig and human lung tissue (fresh snap-frozen) was isolated using the PureLink Mini Kit (Invitrogen) according to the manufacturer’s instructions.

### RNA quantification and quality control

The RNA quantity was established using three methods, one based on the absorbance at 260 nm (A_260_) using spectrometry (Nanodrop) and two kits, Quant-iT RiboGreen RNA Assay Kit (Invitrogen) and Qubit RNA HS Assay Kit (Invitrogen), based on RNA-binding florescent probes. The purity of the samples was assessed by measuring absorbance ratios (A_260_/A_280_ and A_260_/A_230_) using Nanodrop. The RNA integrity was determined by microfluidic capillary electrophoresis using the Agilent RNA 6000 Nano Kit and the 2100 Bioanalyzer Instrument (Agilent).

### Quantitative reverse transcription PCR

Reverse transcription of RNA to cDNA was performed using the T100 Thermal Cycler (Bio-Rad) and iScript RT Supermix (Bio-Rad) with an input of 250 ng RNA (determined by Nanodrop) per sample. Pig *HPRT* and human *RPLP0* were detected using the following primer pairs: AGCCCCAGCGTCGTGATTAG (forward) and AGCAAGCCGTTCAGTCCTGT (reverse) for *HPRT* (Eurofins), and GCGTCCTCGTGGAAGTGAC (forward) and TTCCCCCGGATATGAGGCAG (reverse) for *RPLP0* (Eurofins). For the RT-qPCR analysis, a C1000 Touch Thermal Cycler (Bio-Rad) and SsoAdvanced Universal SYBR Green Supermix (Bio-Rad) (with an annealing temperature of 59 °C). To calculate the PCR efficiency, serial dilutions of cDNA were performed and the cycle threshold value (C_t_) was established for each cDNA concentration. Amplification factor (A) was calculated based on the slope correlating cDNA concentration and C_t_ values; A = 10^1/-S^ (where A = 2 was considered as 100 % PCR efficiency). For all RT-qPCR experiments, two technical replicates were used.

### RNA sequencing

For the library preparation, the TruSeq Stranded mRNA Library Prep Kit (Illumina) was used with an input of 150 ng RNA (determined by the Qubit RNA HS Kit), following the manufacture’s instruction with the exception that 13 cycles instead of 15 were used to enrich DNA fragments. The IDT for Illumina TruSeq RNA UD Indexes (96 indexes) was used to index the different samples for multiplexing. A King Fisher FLEX (Thermo Scientific) was used for automatization of the clean-up steps and a Mastercycler X50s (Eppendorf) was used for the incubations and PCR. The QuantIT dsDNA HS Assay Kit (Invitrogen) was used to determine the DNA concentration of the library. The size and quality of the library was tested using the LabChip GX Touch (Perkin Elmer) with the DNA 1K / 12K / Hi Sensitivity Assay LabChip (Perkin Elmer) and the LabChip DNA High Sensitivity Reagent Kit. A library input of 1.4 pM (with 1 % PhiX Control (Illumina)) was sequenced using the NextSeq 500 instrument (Illumina) with the NextSeq 500/550 High Output v2.5 Kit (Illumina). The FASTQ files created were de-multiplexed using the bclfastq2 software (Illumina) and the software FastQC (Babraham Bioinformatics) was used for the quality control of raw reads. HISAT2 was used for alignment of the reads to the reference genomes and Picard Tools (Broad Institute) was used for quality control (12). All reference genomes were from the Ensembl database (Mouse GRCm38, Pig Ssacrofa11.1, and Human GRCh38) with the GTF annotation (release 99). For the assembly of the alignments into full transcripts and quantification of the expression levels of each gene/transcript, the StringTie software was used (13). The data for this study have been deposited in the European Nucleotide Archive (ENA) at EMBL-EBI under accession number PRJEB39914.

### Bioinformatics

All bioinformatic analyses were done using R packages. Pie charts of transcript type were constructed using total read counts for each RNA type averaged across all samples of the same species. For analyses across species, normalized TPM values were used for genes expressed in at least two of the four samples for each species and in all species. Principle components analysis was done using *prcomp* base package and plotted with *ggbiplot*. Heatmaps were clustered with Euclidian hierarchical clustering on both rows and columns and graphed using the *pheatmap* package.

BisqueRNA was used for bulk RNA-seq deconvolution to estimate cell type proportions using reference-based-deconvolution (14). Biobase was used to generate ExpressionSet objects for bulk and single-cell input data (15). Bulk PCLS data was filtered to include genes expressed in at least 2 of 4 samples for either subject. For comparison to bulk RNA-seq of fresh lung tissue, we combined PCLS data with previously published read counts from GSE99621 and used ComBat_seq for batch correction (16). scRNA-seq data from the IPF cell atlas was used as a reference (17). In brief, raw sparse matrix from GSE136831 was loaded into R using readMM (Matrix package); cell identities, genes, and cell type identities were obtained from GSE136831 metadata (17). The sparse matrix from the single cell reference dataset was downsized as described in the GitHub repository followed by conversion to an ExpressionSet using Biobase. All code is available at https://github.com/lung-bioengineering-regeneration-lab/human_pcls_bisque.

## Results

### RNA yield from pig PCLS using different isolation protocols

The first set of experiments were designed to identify a suitable RNA isolation protocol for PCLS, which generates high yields of RNA from a small amount of tissue. Three protocols for RNA isolation from PCLS were deployed in this study: 1) Qiagen RNeasy Micro Kit (standard protocol), 2) guanidine thiocyanate-phenol (*i.e.* TRIzol) protocol with chloroform RNA phase separation and Qiagen RNeasy MinElute Spin Column clean-up (modified protocol 1), and 3) a modified TRIzol protocol adapted from a previous study by Ogura *et al.* optimizing RNA isolation from immortalized cells suspended in agarose (modified protocol 2) (9). Due to the limited availability of human lung tissue, this first optimization step was performed using PCLS generated from pig lung tissue. To normalize the input of tissue for each isolation, four randomly selected 4 mm diameter (500 µm thickness) PCLS were used for each RNA isolation regardless of protocol. We first assessed RNA yield using absorbance at 260 nm (A_260_) and found that modified protocol 2 led to significantly increased yields as compared to the other protocols with a predicted RNA yield of 1297 ± 125 ng (n = 7) compared to 50 ± 7 (n = 3) and 263 ± 110 (n = 3) ng for the standard protocol and modified protocol 1, respectively (Fig. 1A). As RNA measurement at A_260_ may be affected by contaminants absorbing light at the same wavelength, the RNA yields were additionally measured using two different RNA-binding fluorescent probes, known to be less sensitive for common contaminates. Both of these independent measurements further validated the utility of modified protocol 2 as isolating RNA from small amounts of tissue, however with lower RNA yield measurements compared to that estimated by A_260_ measurements (Fig. 1A).

**Figure 1.**
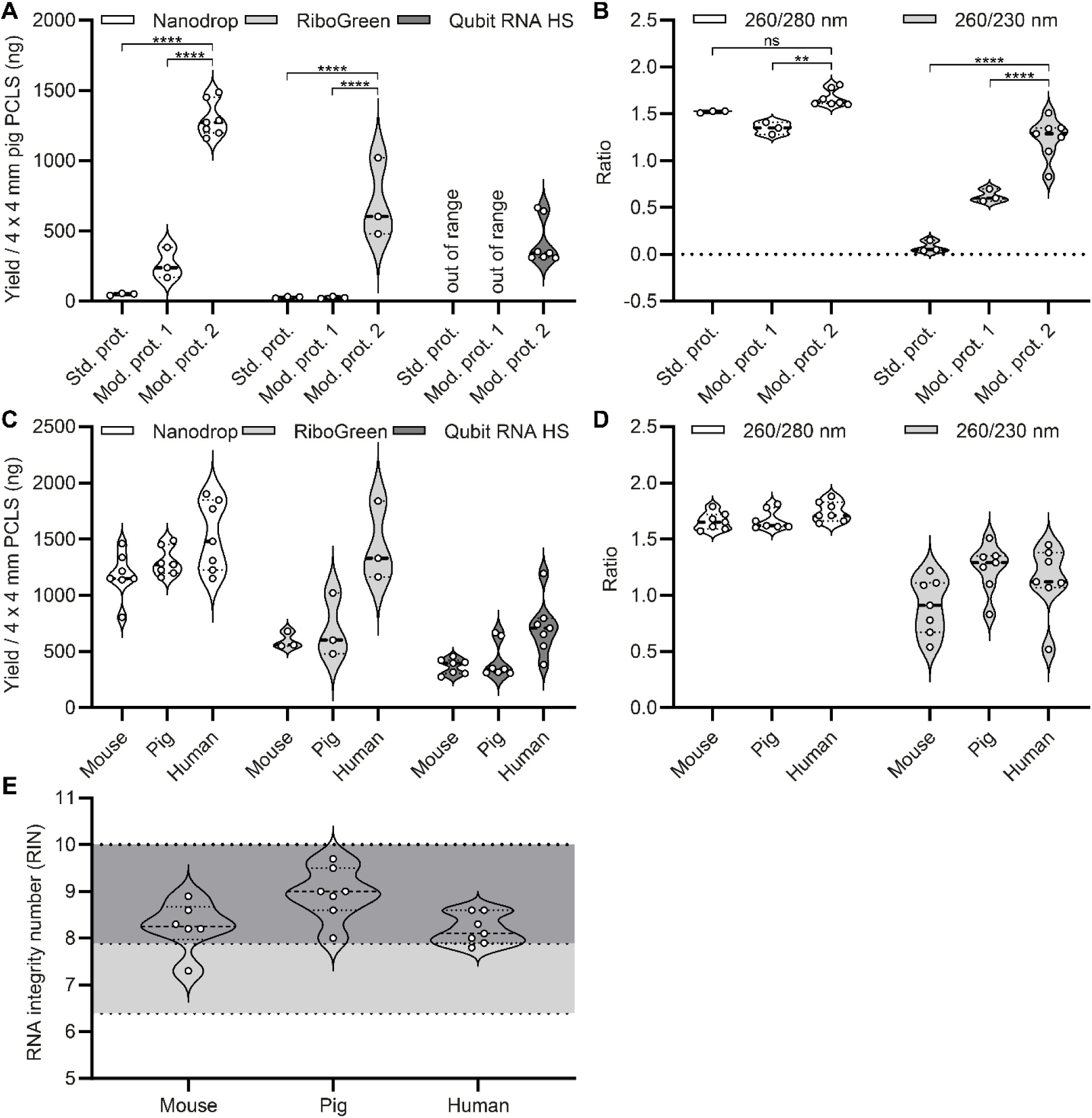
Optimization of RNA isolation from mouse, pig, and human (Patient 1) precision-cut lung slices (PCLS). (**A**) Comparison of the RNA yield from pig PCLS using three different isolation protocols, standard protocol (Std. prot.), modified protocol 1 (Mod. prot. 1), and modified protocol 2 (Mod. prot. 2). Three different RNA quantification methods, based on absorbance at 260 nm (Nanodrop) or RNA-binding fluorescent probes (RiboGreen and Qubit RNA HS), were used. (**B**) Spectrometric assessment of the RNA purity from pig PCLS using the three isolation protocols. (**C-D**) Validation of the yield and purity for modified protocol 2 across species (mouse, pig, and human). The data for pig PCLS is reused from panel A and B as a reference for mouse and human PCLS. (**E**) RNA quality control for mouse, pig, and human PCLS using modified protocol 2, assessed by microfluidic capillary electrophoresis. The light grey area represents RNA integrity number (RIN) values that indicates good quality RNA and the dark grey area represents excellent RNA quality (18). (**A-E**) Each dot represent an independent RNA isolation. To normalize the tissue (PCLS) input in each isolation four 4 mm (diameter) PCLS with a thickness of 500 µm was used for each isolation. To normalize the agarose content, 3 % agarose was used for filling of the lungs of all three species. (**A-B**) 2-way ANOVA with Tukey’s multiple comparisons test (** Adj. P < 0.01, **** Adj. P < 0.0001, ns = not significant).

### Purity of the RNA isolations from pig PCLS

Next, the purity of the RNA isolations was evaluated using absorbance ratios (A_260_/A_280_ and A_260_/A_230_). Optimal values for the A_260_/A_280_ and A_260_/A_230_ ratio for RNA samples is usually considered as 2.0 and 2.1, respectively. Lower ratio values may indicate contaminants such as proteins and phenol for A_260_/A_280_ and carbohydrates, phenol, and guanidine for A_260_/A_230_. However, lower than optimal ratio values are usually accepted for downstream applications (depending on the type of contaminant). Again, modified protocol 2 was superior to the other protocols with ratio values of 1.67 ± 0.09 (A_260_/A_280_) and 1.24 ± 0.22 (A_260_/A_230_) (n = 7) compared to 1.52 ± 0.01 and 1.35 ± 0.07 (A_260_/A_280_) and 0.08 ± 0.06 and 0.62 ± 0.07 (A_260_/A_230_) (n = 3) for the standard protocol and modified protocol 1, respectively (Fig. 1B). As the modified protocol 2 was determined to generate the highest yields with the highest purity values, this protocol was used for isolating RNA in the remainder of this study.

### Yield, purity, and quality of RNA isolated from mouse, pig and human PCLS

PCLS from different species can have different properties (*e.g.* amount of cells in relation to agarose due to the alveolar size and different ECM composition) and may contain different environmental contaminants that potentially could affect the RNA isolation (especially in the case of human lungs). Therefore, the RNA yield and purity were additionally determined for mouse, pig, and human (Patient 1) PCLS. According to yield determination using spectrometry (A_260_), no major differences were observed between species (Fig. 1C). However, fluorescence-based methods indicated slightly higher RNA yields for human PCLS compared to mouse and pig PCLS (P < 0.05) (Fig. 1C). Furthermore, the purities were determined to be similar for the isolations from all three species, except a slightly lower A_260_/A_230_ ratio for mouse *versus* pig PCLS (P < 0.01) (Fig. 1D). These results indicate that modified protocol 2 can achieve high yield and purity across several species. While RNA purity is an important factor for downstream analysis (as contaminants may interfere with *e.g.* enzymatic reactions), another important aspect is RNA degradation (*e.g.* isolation-induced) (18). Thus, the RNA integrity was analyzed using microfluidic capillary electrophoresis. The output from this analysis method, RNA integrity number (RIN), ranks the RNA quality from 1-10 (where a RIN value of 10 indicates non-degraded RNA sample and a RIN of 1 is a totally degraded sample). For downstream analysis using RNA-seq, a RIN value above 6-8 has been suggested as a sample inclusion criteria (18, 19). In this study, the RNA isolations from mouse, pig, and human PCLS were observed to be of similar high quality with RIN mean values of 8.25 ± 0.54 (n = 6), 8.96 ± 0.56 (n = 7), and 8.19 ± 0.32 (n = 7), respectively (Fig. 1E).

### Using RNA isolated from pig and human PCLS for downstream RT-qPCR analysis

Even though modified protocol 2 generated the highest purity values of the protocols tested in this study, both the A_260_/A_280_ and A_260_/A_230_ ratios were below optimal values (*i.e.* 2.0 and 2.1, respectively) (Fig. 1B and 1D). Contaminants could potentially interfere with downstream analysis by for example interfering with different enzymatic reactions. Therefore, the PCR efficiency in RT-qPCR was evaluated by using RNA isolated with modified protocol 2 from pig or human (Patient 1) PCLS or high purity RNA isolated with standard kit protocol from native pig or human (Patient 1) lung tissue (*i.e.* fresh snap-frozen tissue) For this purpose, two housekeeping gene primers were used, pig *HPRT* and human *RPLP0*. We observed similar primer efficiencies for both pig and human PCLS samples as compared to samples from native lung tissue (pig and human PCLS samples were calculated to 104.7 ± 4.7 (n = 3) and 108.4 ± 10.3 (n = 3) %, respectively, with corresponding values of 93.1 and 113.0 % for the native pig and human lung tissue sample, respectively) (Fig. 1A-B). Furthermore, the melt curves displayed no apparent signs of genomic DNA contaminations (Fig. 1C-D). These results taken together indicate that the RNA isolated from PCLS using modified protocol 2 is of sufficient purity for downstream analysis using RT-qPCR and that the contaminants present do not affect the PCR reaction.

**Figure 2.**
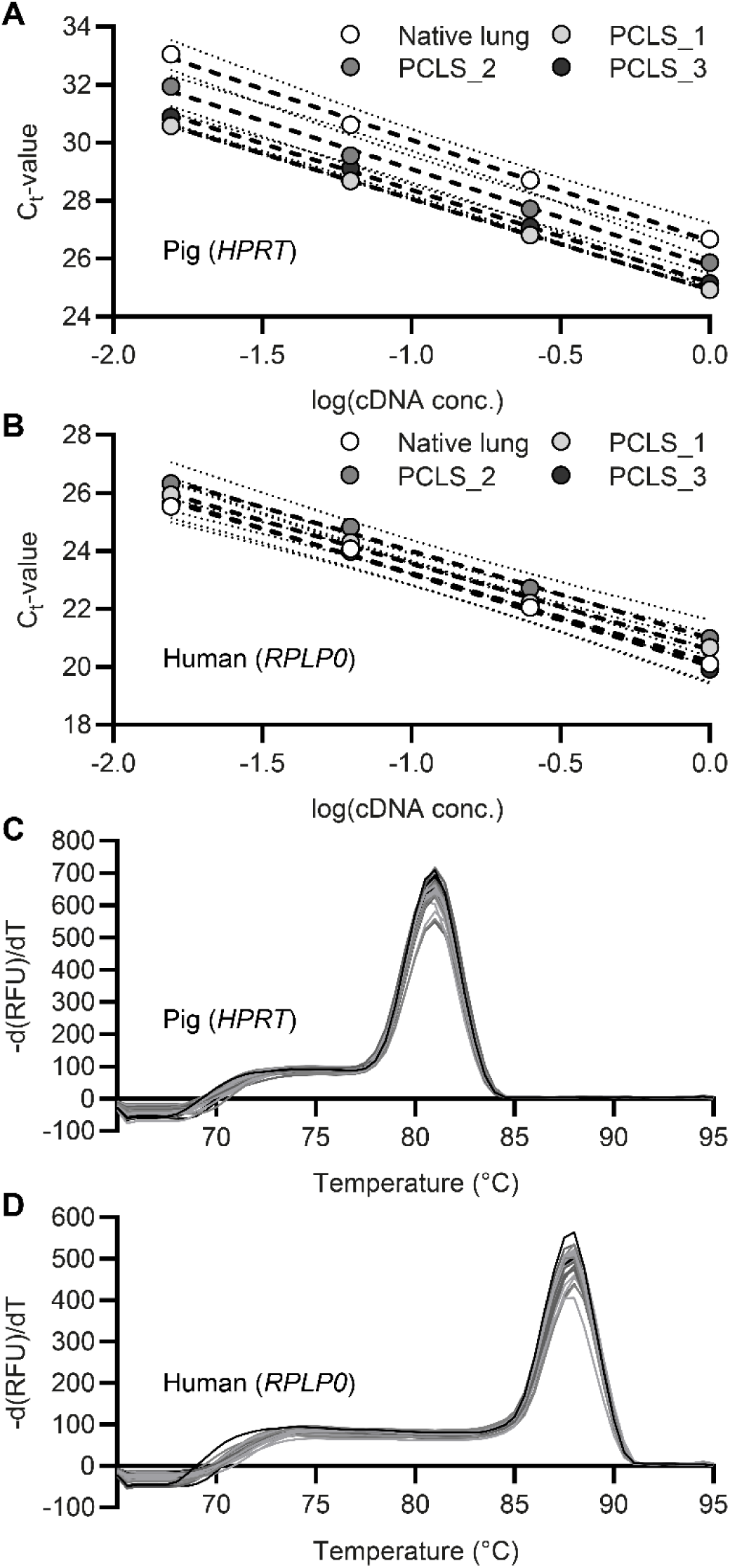
Validation of using RNA, isolated with modified protocol 2, from precision-cut lung slices (PCLS) for downstream analysis with RT-qPCR. (**A**) Comparison of the primer efficiency (*HPRT*) for three independent RNA isolations from pig PCLS using modified protocol 2 and one RNA isolation from native pig lung tissue using standard kit protocol (with high purity, A_260_/A_280_ = 2.07 and A_260_/A_230_ = 2.17). Two technical replicates were used for each data point. (**B**) Comparison of the primer efficiency (*RPLP0*) for three independent RNA isolations from human PCLS (Patient 1) using modified protocol 2 and one RNA isolation from native human lung tissue (Patient 1) using standard kit protocol (with high purity, A_260_/A_280_ = 2.05 and A_260_/A_230_ = 2.05). Two technical replicates were used for each data point. (**C**) The corresponding melt curves for each replicate in panel A. (**D**) The corresponding melt curves for each replicate in panel B.

### Genome-wide RNA sequencing of mouse, pig, and human PCLS

We next sought to determine the possibility of performing RNA-seq using RNA isolated from PCLS with modified protocol 2. Thus, libraries were prepared and sequenced for four independent RNA isolations per species from mouse, pig, and human (Patient 1) PCLS. The mean number of paired-end reads was 26.09 ± 3.30, 27.73 ± 2.46, 27.00 ± 2.37 million for mouse, pig, and human, respectively (Fig. 3A). Of these reads, 80 % or more were aligned only one time, indicating high quality data (Fig. 3A). Of the uniquely aligned reads, the majority of the reads corresponded to protein coding genes (Fig. 3B). However, for mouse and human PCLS, a significant number of reads additionally corresponded to retained introns, nonsense mediated decay, processed transcripts, and long non-coding RNAs (Fig. 3B). For the mouse, pig, and human PCLS samples, the number of genes identified were 15,781, 11,738, 11,924 (expressed in at least 2 out of 4 samples per species), respectively (Fig. 3C). Of these genes, 10,768 genes were common for all three species (Fig. 3C). Principal component analysis and hierarchal clustering, utilizing the commonly expressed genes, displayed clear separation of the samples from the different species (Fig. 3D-E).

**Figure 3.**
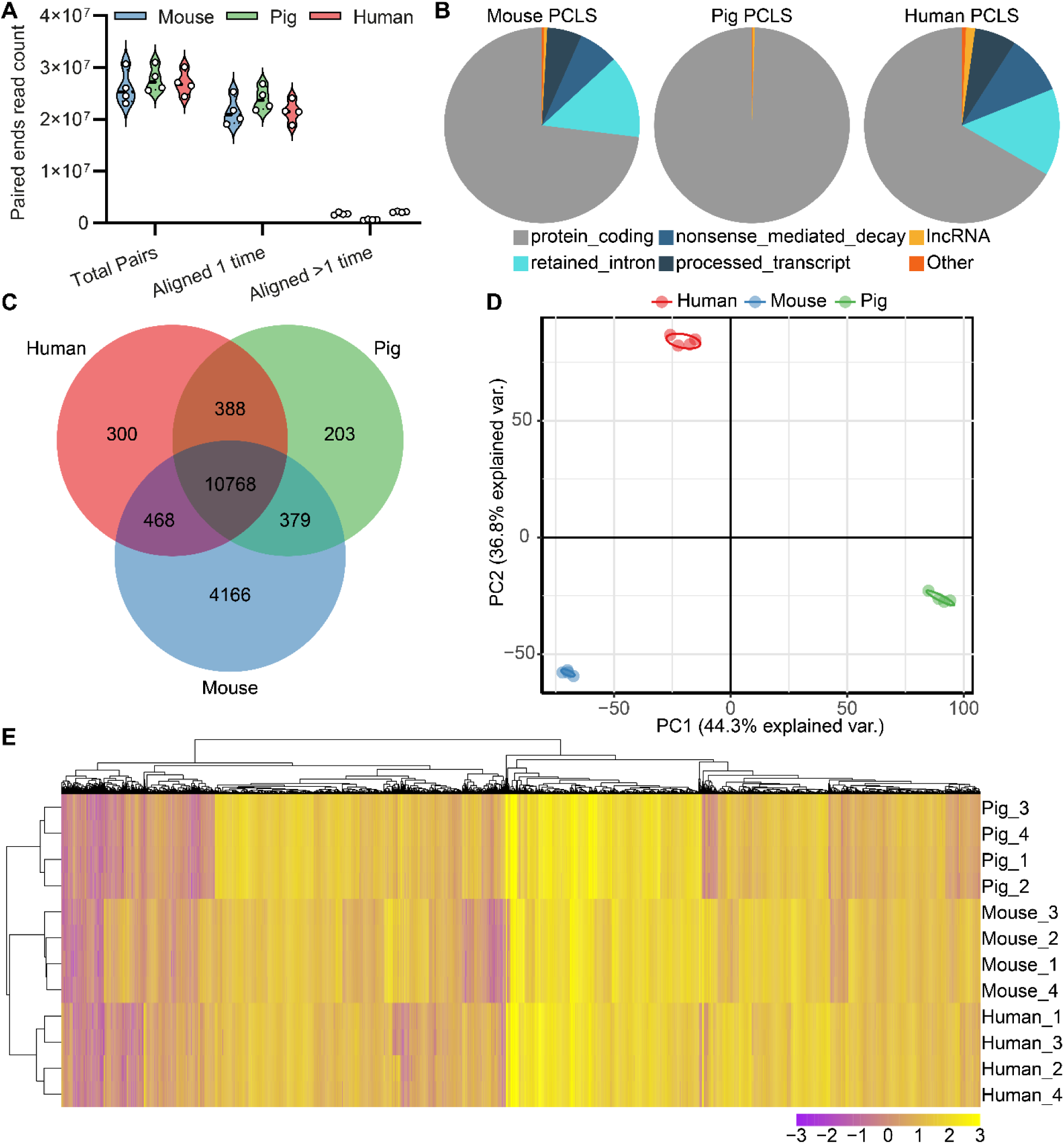
Validation of using RNA, isolated with modified protocol 2, from mouse, pig, and human (Patient 1) precision-cut lung slices (PCLS) for RNA sequencing. (**A**) Number of paired-end reads from mouse, pig, and human PCLS. (**B**) Distribution of uniquely aligned (*i.e.* aligned only one time) reads in regard to gene biotype. (**C**) Commonly expressed genes (present in at least two samples/species) in mouse, pig, and human PCLS. (**D**) Principal component analysis (PCA) of commonly expressed genes (present in at least two samples/species) in mouse, pig, and human PCLS. (**E**) Heat-map of the commonly expressed genes (present in at least two samples/species) in mouse, pig, and human PCLS.

### Deconvolution of human PCLS data predicts the presence of diverse cell types

While a number of cell types have been identified and studied in PCLS (10), this has been restricted to subjective analysis using a limited number of phenotypic markers. Open questions have remained regarding the presence of other cell types, such as immune cells. Therefore, we next sought to assess the cellular composition of human PCLS using Bisque, a recently described deconvolution technique. Bisque has been shown to accurately estimate cell type composition from bulk RNA-seq data using a reference scRNA-seq dataset (7, 14). Here, we used a scRNA-seq reference dataset from the IPF cell atlas, which sequenced and annotated cells present in healthy and diseased lungs (17). We found that Bisque predicted a heterogeneous cell type composition of PCLS, notably including a variety of immune cell types previously not described to be present in PCLS (*e.g.* natural killer cells and dendritic cells) (Fig 4B). Additionally, Bisque predicted changes in cell types known to be associated with IPF (*e.g.* loss of ATII cells in Patient 2 as compared to Patient 1) (20). Furthermore, Bisque predicted the presence of aberrant basaloid cells only in PCLS generated from our IPF patient and not in Patient 1. This matches the recent literature indicating the emergence of this aberrant cell type only in IPF patients (17, 21). We also observed predicted changes in immune cell types known to correspond with lung transplant in Patient 2 (*e.g.* loss of T cells due to cyclosporine, but also increases in cytotoxic T lymphocytes and B cells) (22).

**Figure 4.**
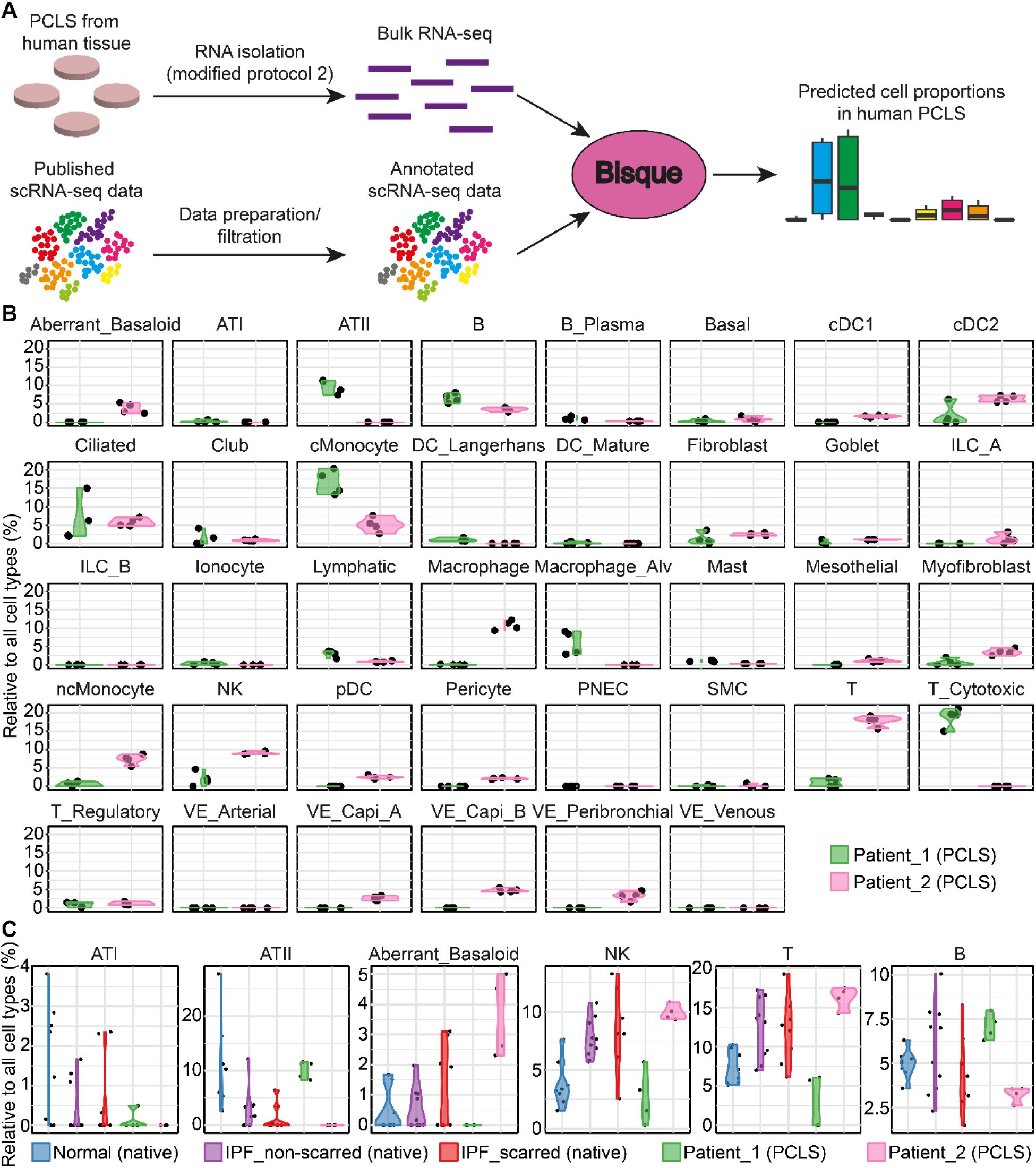
Reference-based deconvolution of human PCLS RNA sequencing (RNA-seq) data using Bisque for prediction of cell type proportions in PCLS. (**A**) Schematic illustration of the workflow. Bulk RNA-seq data was deconvolved using Bisque with the IPF atlas single-cell RNA-seq (scRNA-seq) (GSE136831) as reference (17). (B) Bisque-predicted presence or absence of cell populations in PCLS from Patient 1 (size-mismatch) and Patient 2 (idiopathic pulmonary fibrosis (IPF)). (**C**) Comparison of Bisque-predicted cell types in fresh snap-frozen normal and IPF lung tissue versus PCLS from Patient 1 and 2. For bulk RNA-seq data from fresh snap-frozen lung tissue a publically available data set (GSE99621) was used, containing data for normal, non-scarred IPF, and scarred IPF tissue (23).

### Integration of human PCLS RNA sequencing data with larger cohorts

Having confirmed that our technique results in high quality RNA-seq data amenable to usage with standard and advanced bioinformatic techniques such as supervised clustering and deconvolution, we next sought to explore its usage in tandem with other available cohorts. One major limitation of human PCLS is that human tissue which can be sliced is limited at most centers. Therefore, integrating smaller cohorts of RNA-seq PCLS tissue with larger existing cohorts could be advantageous for placing smaller cohort observations in the context of larger studies. A variety of batch normalization tools exist for comparing RNA-seq datasets generated at different research centers, such as Combat-seq (16). Therefore, we batch normalized the RNA-seq data from our two PCLS patients with RNA-seq data deposited from a cohort comprised of donor lung tissue and IPF tissue from paired scarred and normal looking tissue. Next, we performed Bisque deconvolution to evaluate the cell proportions in our PCLS RNA-seq in comparison to what is present in fresh snap-frozen tissue (23). We found that cell proportions identified in our fibrotic PCLS patient (Patient 2) correlated similarly to fresh tissue from fibrotic regions of patients with fibrosis, as well as cell proportions predicted to exist in the non-scarred regions of patients with fibrosis (Fig 4C). On the other hand, the cellular composition of Patient 1 more closely resembled normal tissue (*e.g.* ATII cell proportion), but also displayed some abnormalities (*e.g.* elevated B cells). (Fig. 4C). As PCLS from Patient 1 were generated from donor tissue explanted two weeks after lung transplantation due to size-mismatch, the altered transcriptome profile from this brief *in vivo* period likely represents the acute but transient immune response to the allograft. Taken together, this confirms that our isolation procedure yields high quality RNA which can be used in advanced computational techniques to produce relevant disease-relevant biological information.

## Discussion

The use of PCLS has recently gained attention as a promising model for studying lung physiology and pathology and as a possible means to screen for novel therapeutics (1). However, the use of PCLS in medical research has been limited by difficulties in obtaining RNA of sufficient quality for non-biased and high through-put analysis methods such as RNA-seq. We here present a novel approach for isolating high quality RNA from small amounts of mouse, pig, and human PCLS, which can be used for downstream RNA-seq and computational analysis.

The high amount of agarose present in the PCLS is considered to be a major source of the difficulty of isolating RNA, which additionally is a well-known problem for other model systems utilizing polysaccharide-rich hydrogels, such as 3D scaffolds for cell culturing (9, 24). To our knowledge, only one study has so far reported feasibility for RNA-seq for rat PCLS (25), but no study has described performing RNA-seq from PCLS generated from human tissue. The previous study, utilizing rat tissue, (25), used a low agarose concentration (0.75 %) for filling the rat lungs, which most likely facilitated the isolation of high quality RNA. However, the usage of an equivalent percentage of agarose is usually not a possibility for generating PCLS from larger species, such as pig or human (especially from diseased lung explants), where an agarose concentration of 3 % is most commonly used (1). Additionally, a recent study by Niehof *et al.* reported improved RNA isolation from human PCLS containing 1.5 % agarose for RNA microarray analysis, using a magnetic bead-based isolation approach (8). However, they did not validate their protocol for RNA-seq or for PCLS containing agarose concentrations above 1.5 %. In the current study, we therefore aimed to develop a reliable method for isolating high yield, purity, and quality RNA from human PCLS containing up to 3 % agarose, suitable for RNA-seq. We additionally wanted the method to be reproducible for several species, quick to use, and free of steps requiring specialized and expensive equipment.

We performed the first optimization experiments on PCLS derived from pig tissue and confirmed that the standard RNA isolation protocol used previously, by us and others (25), suffers from low RNA yields and purity. Next, we tried guanidine isothiocyanate-phenol mixtures (*e.g.* TRIzol) due to the fact that it has long been used for extracting RNA from polysaccharide-rich samples (26). However, we found that TRIzol, together with phase separation using chloroform (where RNA is retained in the aqueous phase) only slightly improved the yield, thus implying that the agarose was still interfering with proper RNA isolation. In line with this, recent results by Ogura *et al.* suggest that the TRIzol-dissolved agarose (used as a 3D scaffold for cell culturing) is retained in the aqueous phase together with the RNA (9). This finding could explain the low yield using chloroform-induced phase separation. Therefore, we next modified the protocol by excluding the phase separation step and found that the yield was significantly increased, likely by maintaining the agarose in solution during the initial steps of the spin column RNA purification. However, as chloroform-induced phase separation is an effective way of excluding unwanted isolation of genomic DNA, we subsequently included a step for DNA removal (*e.g.* DNAase) and found that it was critical for successful implementation of the protocol (data not shown). In addition to improving the yield and quality of mRNA for downstream analysis using RT-qPCR and RNA-seq, we also observed that the yield of micro-RNA (miRNA) was increased using modified protocol 2 compared to commercial kits for miRNA isolation (data not shown).

The highest purity values were obtained using modified protocol 2. However, the optimized protocol still generated sub-optimal spectrometry ratio values (especially for A_260_/A_230_), thus indicating some remaining contaminants. Additionally, the lower RNA yield detected using RNA-binding florescent probes, which are less sensitive for contaminants compared to spectrometry, further validated the presence of contaminants. Nevertheless, the contaminants observed in this study were not found to interfere with downstream RT-qPCR analysis which can be highly sensitive to contaminant inhibiting PCR-related enzymatic reactions. This finding is in accordance with the previous study by Niehof *et al.*, who reported similar purity values for RNA isolated from human PCLS (with 1.5 % agarose) without any interference of RT-qPCR (8).

Using RNA isolated from PCLS with modified protocol 2 for RNA-seq generated high quality data as indicated by the quality control of raw reads and read alignment (FastQC and Picard Tools, data not shown). For example, the ratio of uniquely mapped reads to the total number of reads were high (≥ 80 %), median insert sizes around 150 bp (data not shown), and the mapped read GC content aligned well with the reference genome (data not shown), together indicating data of high quality (6). The major biotype identified in our data was for protein coding transcripts, whereas biotypes such as ribosomal RNA and short RNAs constituted only minor fractions, as expected from the library preparation used in this study. The much higher fraction of protein coding transcripts for the pig samples compared to mouse and human is most likely due to the lower degree of annotation for the pig genome. Principal component analysis and hierarchal clustering of commonly expressed genes for mouse, pig, and human PCLS resulted in clear clustering of the samples, which is in line with a recent study comparing native lung tissue from similar species using scRNA-seq (27). Importantly for this study, however, tight intra-species clustering indicate consistent gene expression patterns in each of the four independent samples from each species.

Deconvolution of bulk RNA-seq has recently emerged as a powerful technique to use scRNA-seq data with bulk RNA-seq to predict the cell composition of bulk RNA. Single-cell sequencing offers unprecedented insight into changes in cellular composition of disease, but several major drawbacks remain. All of the published human lung scRNA-seq datasets are heavily immune cell biased, which skews predictions using current deconvolution tools. For example, macrophages (resident or alveolar) can constitute around 59 % of entire datasets (17). In order to overcome this limitation and attempt to more accurately predict cellular composition in our bulk RNA, we computationally reduced the number of macrophages for each patient to similar proportions found in recently published mouse lung scRNA-seq datasets (28). While this allowed us to predict the presence of other cell types in the PCLS, the current predictions are likely more indicative of the presence or absence of cell types and their relative presence between conditions and not representative of true abundance. Single-nuclei RNA-seq is known to be less biased than single-cell RNA-seq in this regard and the generation of these datasets and may help overcome this limitation for use in deconvolution of bulk RNA-seq data in the future. Alternatively, techniques to separate cells or nuclei from the agarose in PCLS could be used. However, single-cell or - nuclei RNA-seq remain costly and deconvolution offers an opportunity to first screen multiple conditions to identify conditions of interest on which to perform single-cell or -nuclei RNA-seq. Additionally, deconvolution techniques are also known to predict slightly different cell compositions based on the phenotypic marker list used and the number of patients present in the scRNA-seq cohort. Furthermore, the presence, viability, and function of all of these predicted cell types needs to be confirmed at the cellular resolution in PCLS using immunofluorescence of multiple phenotypic markers or spatial transcriptomics/*in situ* hybridization techniques. Despite these limitations, the prediction of the presence of diverse immune cell types from our bulk RNA-seq data is encouraging for use of PCLS in additional areas of respiratory research, including precision medicine approaches.

This new method of isolating RNA from PCLS has the possibility of significantly accelerating the use of PCLS for pulmonary research and drug discovery. Furthermore, this new RNA isolation method may prove useful in other research fields, such as tissue engineering and 3D cell culture systems, which struggle with isolating RNA from polysaccharide-rich hydrogels.

## Acknowledgments

The authors thank all the members of the Lung Bioengineering and Regeneration (LBR) Laboratory (Department of Experimental Medical Sciences, Lund University) for helpful discussions throughout the project and Prof. Richard Ingemansson (Department of Thoracic Surgery, Lund University) for kindly supplying the IPF tissue and valuable clinical insights. We are grateful to Tomas Jansson (Department of Biology, Lund University) for performing the RNA quality control. We thank the Center for Translational Genomics (Lund University) and Clinical Genomics Lund (SciLifeLab) for providing sequencing service. The Knut and Alice Wallenberg foundation is acknowledged for generous support (DEW, NDL, and SLI) and the Swedish Research Council (DEW, Registration Number 2018-02352). Further support is acknowledged from an American Thoracic Society Foundation Unrestricted Grant (DEW), an ERASMUS+ award and short-term Fellowship from the European Respiratory Society (WL) and the Crafoord Foundation (JS, Reference Number 20190708).

